# The role of replication clamp-loader protein HolC of *Escherichia coli i*n overcoming replication / transcription conflicts

**DOI:** 10.1101/2020.12.02.408393

**Authors:** Deani L. Cooper, Taku Harada, Samia Tamazi, Alexander E. Ferrazzoli, Susan T. Lovett

**Author notes:** Alexander Ferrazzoli passed away on August 4, 2020. Karp Family Research Laboratories, 1 Blackfan Cir, Boston, MA 02115.

## Abstract

In *Escherichia coli,* DNA replication is catalyzed by an assembly of proteins, the DNA polymerase III holoenzyme. This complex includes the polymerase and proofreading subunits as well as the processivity clamp and clamp loader complex. The *holC* gene encodes an accessory protein (known as x) to the core clamp loader complex and is the only protein of the holoenzyme that binds to single-strand DNA binding protein, SSB. HolC is not essential for viability although mutants show growth impairment, genetic instability and sensitivity to DNA damaging agents. In this study, to elucidate the role of HolC in replication, we isolate spontaneous suppressor mutants in a *holCΔ* strain and identify these by whole genome sequencing. Some suppressors are alleles of RNA polymerase, suggesting that transcription is problematic for *holC* mutant strains or *sspA,* stringent starvation protein. Using a conditional *holC* plasmid, we examine factors affecting transcription elongation and termination for synergistic or suppressive effects on *holC* mutant phenotypes. Alleles of RpoA (α), RpoB (β) and RpoC (β’) RNA polymerase holoenzyme can partially suppress loss of HolC. In contrast, mutations in transcription factors DksA and NusA enhanced the inviability of *holC* mutants. Mfd had no effect nor did elongation factors GreA and GreB. HolC mutants showed enhanced sensitivity to bicyclomycin, a specific inhibitor of Rho-dependent termination. Bicyclomycin also reverses suppression of *holC* by *rpoA rpoC* and *sspA.*These results are consistent with the hypothesis that transcription complexes block replication in *holC* mutants and Rho-dependent transcriptional termination and DksA function are particularly important to sustain viability and chromosome integrity.

**IMPORTANCE:** Transcription elongation complexes present an impediment to DNA replication. We provide evidence that one component of the replication clamp loader complex, HolC, of *E. coli* is required to overcome these blocks. This genetic study of transcription factor effects on *holC* growth defects implicates Rho-dependent transcriptional termination and DksA function as critical. It also implicates, for the first time, a role of SspA, stringent starvation protein, in avoidance or tolerance of replication/replication conflicts. We speculate that HolC helps resolve codirectional collisions between replication and transcription complexes, which become toxic in HolC’s absence.

## INTRODUCTION

The ability to replicate DNA faithfully is critical for the survival of all organisms. The replication fork very frequently encounters barriers that need to be overcome to ensure cell survival and genetic stability (Cox et al. 2000; Kowalczykowski 2000). Such barriers may be breaks, nicks, or modified bases in the DNA template, damage to the dNTP pool or nascent strand, tightly bound proteins, transcription complexes, and DNA secondary structures. Single-stranded gaps left behind by the fork can be filled by a number of mechanisms found broadly across organisms, including homologous recombination with the sister chromosome, translesion DNA synthesis and template-switching (Lovett 2017).

The bulk of DNA replication in the bacterium *Escherichia coli* is catalyzed by the DNA polymerase III holoenzyme (McHenry 1991; O’Donnell 2006). This multi-subunit complex consists of the core DNA polymerase assembly with a proofreading exonuclease subunit (α,ε,φ)^1^ the processivity clamp (β) and an associated clamp-loader complex ([τ/γ]_3_δδ’) with its accessory complex (χψ). The processivity clamp is a ring-like structure that encircles DNA and tethers DNA polymerases to their templates, conferring processivity to DNA synthesis. The pentameric clamp loader complex can both load and unload the clamp, a cycle that must be completed each round of Okazaki fragment synthesis on the lagging strand. The structure of the clamp and the clamp loader are conserved in all domains of life; in archaea and in eukaryotes, they are known as PCNA (proliferating nuclear antigen) and RFC (replication factor C), respectively. In E*. coli,* the clamp binds all of its 5 DNA polymerases (Dalrymple et al. 2001); in addition to DNA pol III, it binds pol I, involved in Okazaki fragment maturation and RNA primer processing (Okazaki et al. 1971), and the DNA repair polymerases II, IV and V (Napolitano et al. 2000).

Most of the proteins in the DNA polymerase III holoenzyme are essential for viability with some notable exceptions, two of which are HolC/HolD (or χ/φ, respectively) that form an accessory heterodimer that binds to the core pentameric clamp loader complex. HolC and HolD are not ubiquitous in bacteria and are found only in γ-proteobacteria, although there may be more unrelated proteins that play similar roles in other bacteria. HolC is of particular interest because it is the only protein of the DNA pol III holoenzyme that binds single-strand DNA binding protein, SSB, at a site distinct from its interaction with HolD (Marceau et al. 2011). At the opposite face of its interaction site with HolC, HolD interacts with the DnaX-encoded subunits of the pentameric clamp loader (Gulbis et al. 2004). Therefore, together HolC and HolD form a bridge between SSB-coated template DNA, the pentameric clamp loader complex and the rest of the DNA pol III holoenzyme.

In vitro studies have suggested a number of roles for the HolC/D accessory complex in DNA replication. There is evidence that the HolC/D complex assists assembly and stability of the clamp loader complex (Olson et al. 1995) and increases its efficiency of clamp loading (Anderson et al. 2007). HolC, through its interaction with SSB, aids the engagement of DNA pol III with RNA primers and generally stabilizes interaction of the replisome with its template (Glover and McHenry 1998; Kelman et al. 1998; Marceau et al. 2011). HolD, through its interaction with DnaX proteins, induces higher affinity of the clamp loader for the clamp and for DNA (Xiao et al. 1993; Gao and McHenry 2001).

Deletion mutants of HolC are viable but grow quite poorly and their cultures rapidly develop genetic suppressor variants. HolC mutants, even when grown under conditions that ameliorate their inviability, exhibit elevated rates of local genetic rearrangements, as do many mutants with other impairments in the DNA replication machinery (Saveson and Lovett 1997). Mutants of HolC lacking its interaction with SSB causes temperature-dependent induction of the SOS DNA damage response and cell filamentation, with a block to chromosome partitioning (Marceau et al. 2011). All in all, these phenotypes point to the aberrant nature of replication in the absence of HolC function.

Michel and collaborators have done several studies of suppressor mutations that improve the viability of strains that lack HolC’s partner, HolD. A duplication of the *ssb* gene is one such suppressor, which suppresses loss of either HolC or HolD or both (Duigou et al. 2014). This suggests that ssDNA gaps accumulate in HolCD mutant strains; extra SSB may protect ssDNA and recruit repair factors (Shereda et al. 2008) to aid gap-filling. Accumulation of ssDNA induces *E. coli’s* DNA damage response, the “SOS” pathway; blocking this with a non-inducible allele of the SOS repressor, LexA (LexAind^-^), also improves the viability of HolD mutants (Viguera et al. 2003). The negative effect of the SOS response in HolD mutants stains is due to increased expression of the translesion DNA polymerases, DNA polymerase II and DinB (Viguera et al. 2003) and, to a lesser extent, to a SulA-dependent block to cell division. Mutations in the replisome-associated ATPase, RarA (Barre et al. 2001), implicated in DNA polymerase exchange (Shibata et al. 2005), are also partial suppressors and its suppression of HolD is epistatic to LexA(Ind^-^)(Michel and Sinha 2017), indicating a common mechanism. These results suggest that the accumulation of replication gaps in HolD mutants triggers the SOS response, including the up-regulation of translesion DNA polymerases pol II and pol IV who compete with DNA pol III, replacing it on the clamp. Because these polymerases are slower or more error-prone than pol III (Qiu and Goodman 1997; Wagner et al. 1999), this polymerase exchange may be deleterious. An L32V allele of the clamp-loader subunit, DnaX, to which HolD binds, was also found as a suppressor (Michel and Sinha 2017), and may increase the stability or functionality of the clamp loader complex in the absence of HolD. Likewise, mutations affecting K^+^ import, TrkA and RfaP may also suppress HolD by this mechanism (Durand et al. 2016). Finally, inactivation of the stringent starvation protein, SspA, suppresses HolD by an unknown mechanism, genetically distinct from SOS, RarA and TrkA (Michel and Sinha 2017).

It had been assumed that the function of HolC and HolD are obligately linked. However, HolC is implicated in repair of damaged forks in a way that HolD is not. HolC physically interacts with a putative DNA helicase of the XP-D/DinG family, YoaA, that is induced by DNA damage; both enhance survival to the replication chain terminator nucleoside 3’ azidothymidine, AZT, (Brown et al. 2015) that produces gaps during replication (Cooper and Lovett 2010). In unpublished work, we have provided evidence that the HolC YoaA and HolC HolD complexes are mutually exclusive. Both HolD and YoaA appear to bind to the same surface of HolC including residues W57 and P64, at a site distal to the residues required for interaction with SSB (Gulbis et al. 2004). We proposed that, after DNA damage and the accumulation of unreplicated DNA, this mechanism allows the recruitment of the YoaA helicase to the fork, without accompanying DNA pol III molecules.

To clarify the role of HolC in replication and repair, we characterize here a number of spontaneous suppression mutations to *holC*.

## RESULTS

A *holCΔ* strain was grown overnight in minimal medium at 30°, conditions that minimize toxicity and plated on LB or LB with AZT and incubated at 37° overnight. Under these conditions, *holCΔ* strains grown poorly and form small colonies. We isolated large colony variants, which were purified to single colonies on minimal medium at 30°. Because AZT must be phosphorylated before it can be incorporated into DNA, many spontaneous AZT-resistant derivatives have mutations in the *tdk* gene, encoding thymidine kinase (Elwell et al. 1987) of the thymidine salvage pathway, and often are deletions of all or part of the locus. We used a colony PCR assay to screen these out. 26 strains, 11 derived from selection on LB and 15 from selection on LB-AZT, with a wt-length *tdk* locus were frozen and DNA prepared for whole genome sequencing. Of the 15 AZT-resistant isolates, 6 had point mutations in the *tdk* gene and were not pursued further. Among the remaining strains several piqued our interest: 3 isolates had mutations in RNA polymerase (RpoA R191C, RpoA dup[aa179-186] and RpoC E756K), one had an alteration in the replication fork helicase, (DnaB E360V,) and 2 had mutations in stringent starvation protein, SspA (SspA Y186S and A to C in its upstream Shine-Dalgarno sequence). All of these except RpoA R191C were isolated as faster growing variants on LB; RpoA R191C was selected as an AZT-resistant isolate.

To study the genetic properties of these suppressors in the absence of selection for growth, we engineered a *holC* conditional mutant, with *holC* deleted on the chromosome and a plasmid encoding *holC+* that can be retained only in the presence of IPTG. In medium with IPTG, cells are HolC^+^; without IPTG, the complementing plasmid is lost and the *holC* mutant phenotype is revealed.

In a study by Michel and colleagues (Michel and Sinha 2017), loss-of-function mutations in *sspA,* a transcriptional activator protein, were found to suppress *holD.* We recovered two alleles of *sspA* among our suppressed *holC* strains. Because it is not clear what effects these alleles would have on *sspA* function, rather than characterizing them further, we assayed the consequence of an *sspA* knockout mutation on *holC* phenotypes in the conditional strain. Growth defects of *holC* mutants (lacking the pAM-*holC* plasmid) were enhanced with richer growth medium (LB > min CAA > min) and at higher temperatures. Concomitant loss of *sspA* function provided full suppression of *holC* growth defects in all conditions (Figure 1). Suppression of *holC* by *sspA* was most dramatic under the most restrictive condition, LB at 42°, where plating efficiency was increased by 4 orders of magnitude. Mutants in *<u>holC</u>* cured of the complementing plasmid showed a broad distribution of increased cell lengths (Figure 2), including long cell filaments; addition of *sspA* largely returned this distribution to one similar to wt.

**Figure 1:**
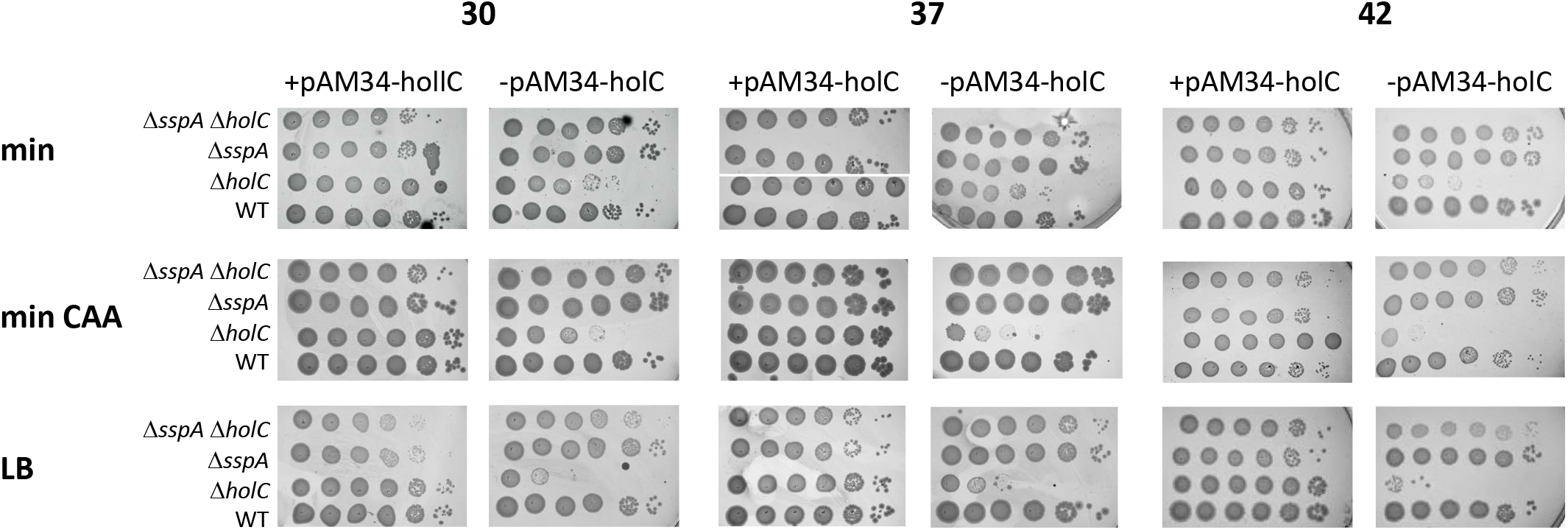
Suppression of *holC* by *sspA*. 10-fold serial dilutions of cultures with and without the *holC* complementing plasmid (pAM34-*holC*) were plated on minimal glucose (min), minimal glucose casamino acids (min CAA) or LB media and incubated at the indicated temperature.

**Figure 2.**
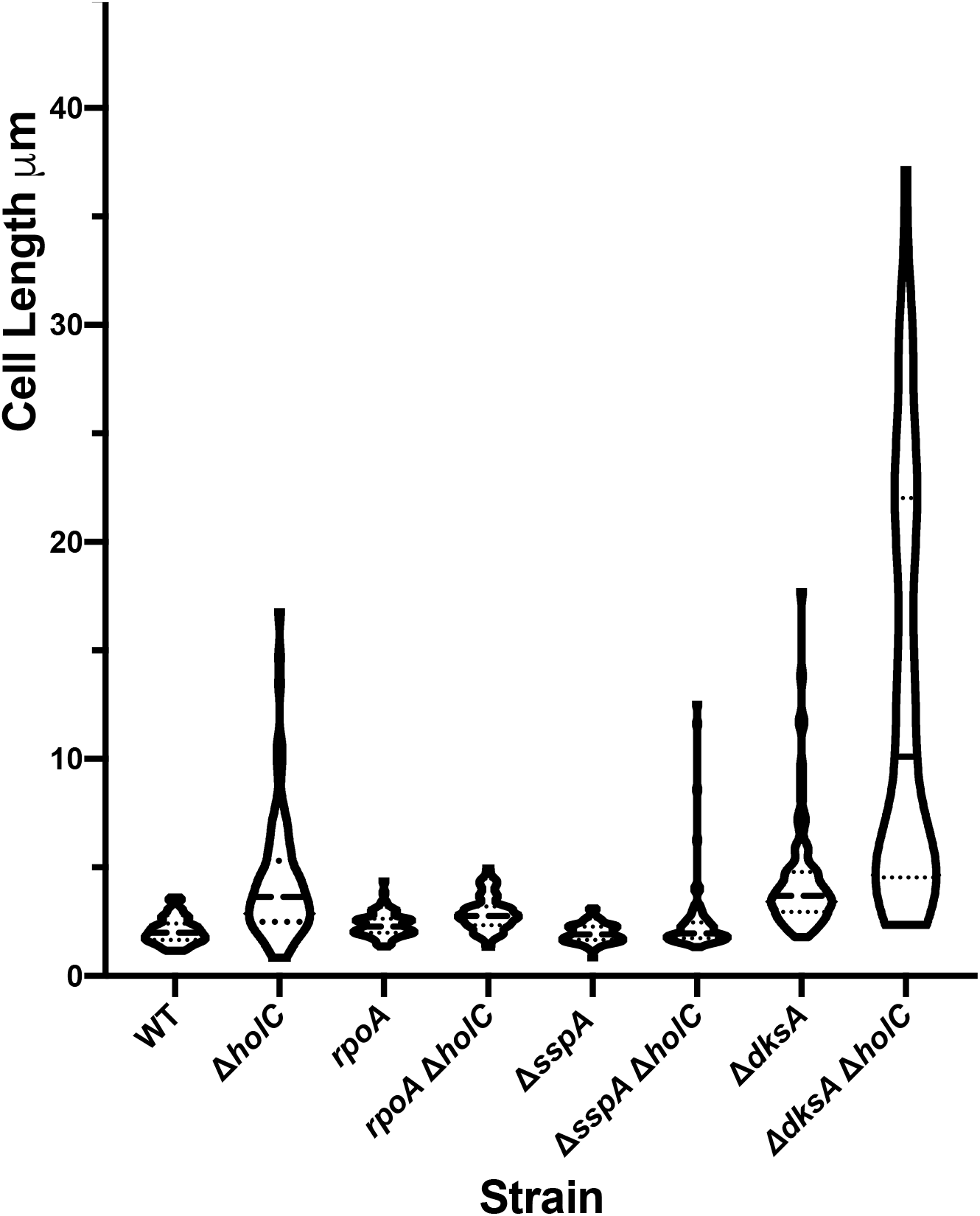
Violin plot of the cell length distribution of *holC* mutant derivatives grown in min CAA at 30° as determined by microscopy. The long dashed line indicates the median value and dotted lines the quartile values.

Michel et al. also found that in *holD* mutants, RecF-dependent induction of the SOS response contributes to its poor growth phenotype (Viguera et al. 2003). Some toxicity conferred by *holD* could be relieved by inactivation of SOS-induced DNA polymerases, pol II *(polB)* and pol IV *(dinB),* implicating polymerase exchange as contributing to toxicity in *holD* strains. We likewise found a modest increase in plating efficiency of the *holC* mutants in strains lacking *polB* or *dinB* (Figure 3, suppression is most evident at 30 and 37°). As observed for *holD* (Viguera et al. 2003), we saw little or no suppression of the growth defects of *holC* by *sulA,* the cell division inhibitor induced by the SOS response.

**Figure 3.**
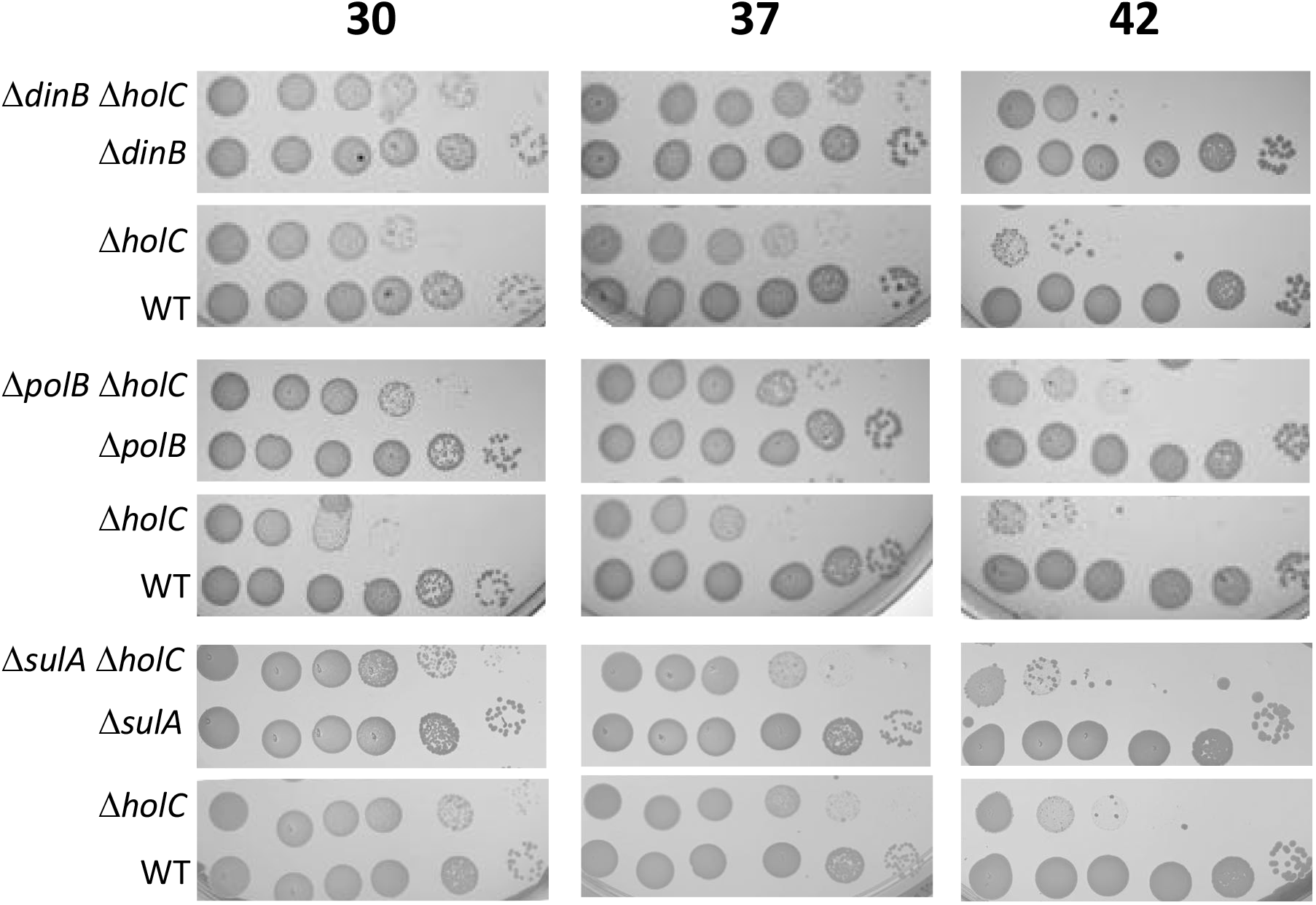
Suppression of *holC* by SOS-induced functions. 10-fold serial dilutions of cultures cured for the *holC* complementing plasmid (pAM34-*holC*) were plated on minimal glucose (min) medium and incubated at the indicated temperature.

Most intriguing were the suppressor isolates affecting RNA polymerase (RNAP), which were not identified in prior studies of *holD* suppression. By genetic backcrosses, we showed that the RpoA duplication of amino acids 179-186 was sufficient to suppress the the poor growth of *holC* mutants (Figure 4) under many conditions. Growth of the *holC* mutant in the absence of IPTG was poor, especially on rich medium and at higher temperatures. The suppression by RpoAdup[aa179-186] of *holC* was not complete and some inviability is retained at higher temperatures and on LB (Figure 4). However, the RpoAdup[aa 179-186] single mutant strain itself was LB-sensitive and temperature-sensitive. In addition, the *holC^+^* plasmid (a ColEI, medium copy derivative) appeared to be toxic to *rpoA* mutants, especially on LB and at higher temperature: survival was enhanced after plasmid loss (right panel vs. left panel, Figure 4). Filamentation to larger cell length in *holC* strains was also ameliorated by RpoAdup[aa179-186] (Figure 2).

**Figure 4.**
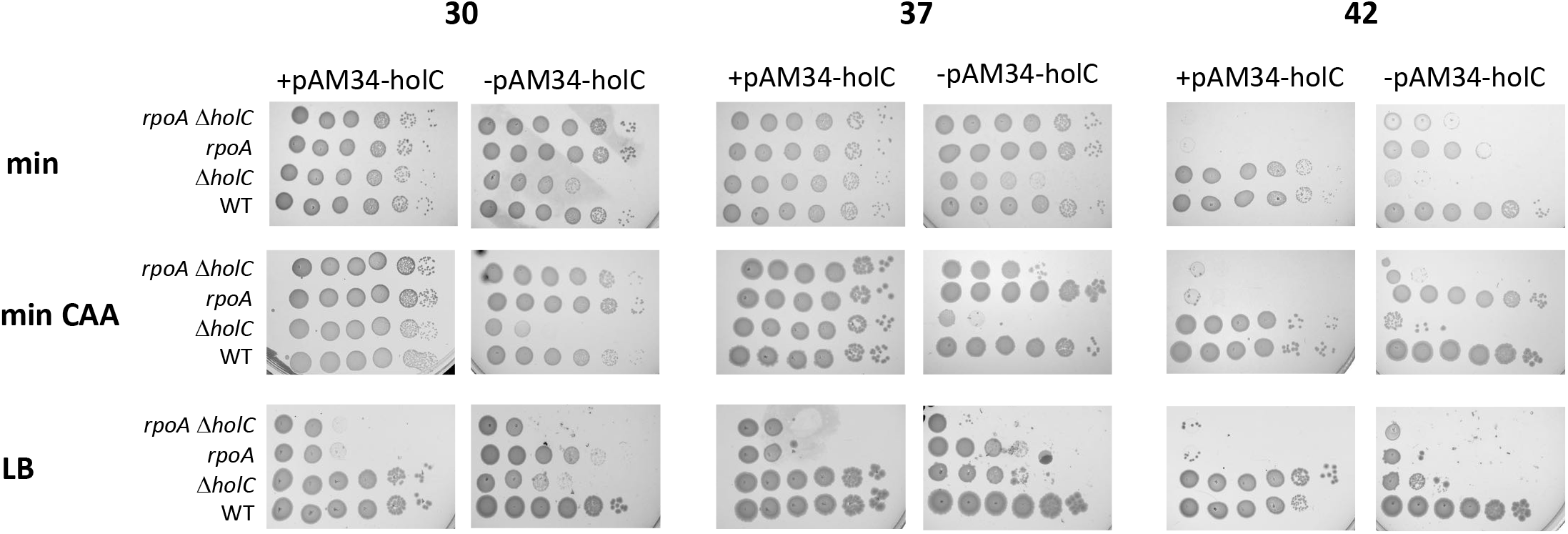
Suppression of *holC* by *rpoA* dup[aa 179-186]. 10-fold serial dilutions of cultures with and without the *holC* complementing plasmid (pAM34-*holC*) were plated on minimal glucose (min), minimal glucose casamino acids (min CAA) or LB media and incubated at the indicated temperature.

The other *rpoA* allele isolated in the screen, R191C, is the same mutation as *rpoA101,* a well-characterized temperature-sensitive allele of RNAP α (Kawakami and Ishihama 1980; Igarashi et al. 1990); RNAP assembles with normal kinetics in this mutant but is unstable, with β and β’ rapidly degraded. Because its intrinsic temperature-sensitivity would confound that of *holC,* we did not characterize this allele further.

We were unable to recover the *rpoC* E756K strain, but during the course of genetic analysis, we discovered that a *rpoC*::GFP fusion allele was able to partially suppress *holC* phenotypes (Figure 5). This allele was considered to be functional but somewhat temperature-sensitive (,Cabrera and Jin 2003) although it can sustain viability in the absence of other *rpoC* genes in lower temperatures. This suppressive effect confirms that it must be perturbed in some way. Like *rpoBdup[aa* 179-186] *holC^+^* strains, *rpoC*::GFP *holC*^+^strains were also LB-sensitive, especially at high temperature.

**Figure 5.**
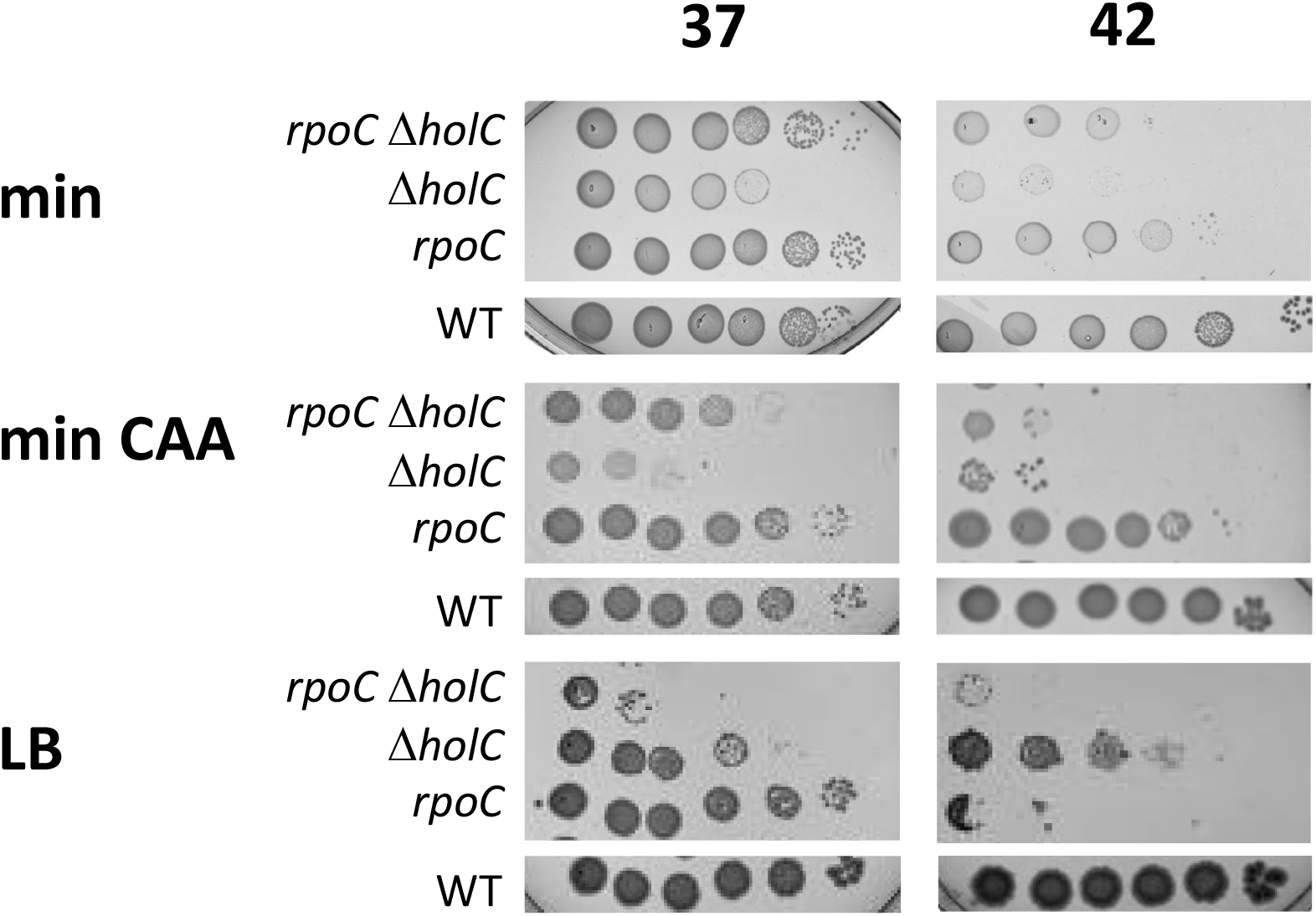
Suppression of *holC* by *rpoC*::GFP. 10-fold serial dilutions of cultures without the *holC* complementing plasmid were plated on minimal glucose (min), minimal glucose casamino acids (min CAA) or LB media and incubated at the indicated temperature.

Transcription complexes pose a major impediment to the replication fork ((Mirkin and Mirkin 2007; Merrikh et al. 2012; Aguilera and Gaillard 2014). To decipher the mechanism of RNAP-suppression of a DNA replication mutant, we examined mutants in several factors known to modulate transcription elongation for their effects, positive or negative, on *holC* mutant phenotypes. GreA and GreB are elongation factors that reactivate backtracked transcription elongation complexes by promoting cleavage of the RNA 3’ terminus to reposition it in the active center of the enzyme (Borukhov et al. 1992; Orlova et al. 1995). Neither of these functions are essential for viability and neither had effects on *holC* phenotypes (Supplemental Figure S1 and data not shown). Mfd mediates transcription-coupled repair, where excision repair proteins are recruited to sites of RNAP arrest (Selby and Sancar 1993). In addition, through its ATP-dependent translocase activity Mfd promotes RNAP release from the DNA template (Park et al. 2002; Deaconescu et al. 2006). Loss of Mfd neither enhanced nor suppressed *holC* inviability (Supplemental Figure 1).

DksA is structurally similar to the Gre proteins and binds to the secondary channel of RNAP; it affects both the initiation and elongation properties of RNAP especially in the presence of the signaling molecule ppGpp (reviewed in (Gourse et al. 2018)). In vivo, there is evidence that DksA alleviates conflicts between replication and transcription, preventing replication arrest by stalled transcription complexes during amino acid starvation (Tehranchi et al. 2010). Mutants in *dksA* had synthetic growth defects when combined with *holC* (Figure 6), indicating that DksA function protects replication in the absence of *holC*. Loss of *dksA* also exacerbated cell filamentation in *holC* mutants (Figure 2). Like DksA, a mutant of NusA, *nusA11,* reduced the plating efficiency of *holC* mutants (Figure 7). NusA potentiates Rho-dependent termination, which in vivo prevents replication fork collapse and double-strand break formation (Washburn and Gottesman 2011).

**Figure 6.**
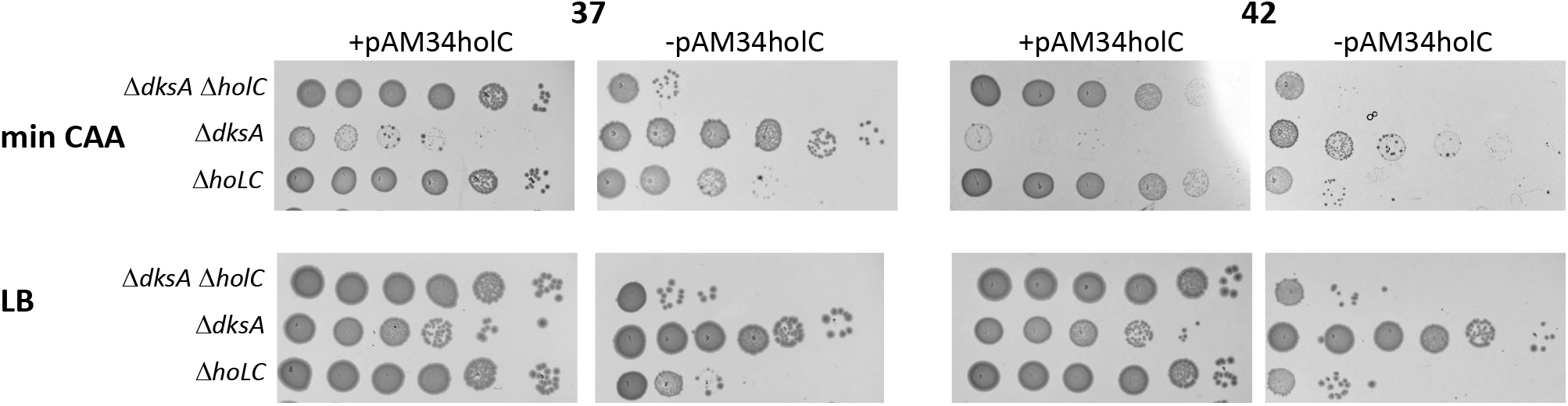
Enhancement of *holC* growth defects by *dksA*. 10-fold serial dilutions of cultures with and without the *holC* complementing plasmid (pAM34-*holC*) were plated on minimal glucose casamino acids (min CAA) or LB media and incubated at the indicated temperature.

**Figure 7.**
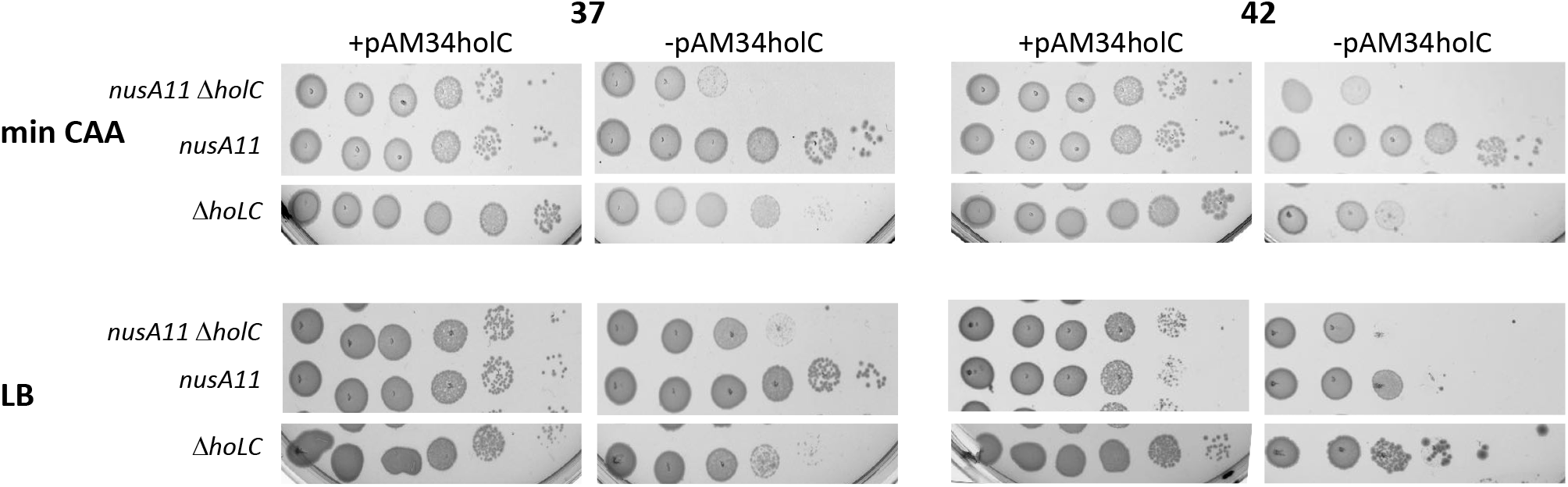
Enhancement of *holC* growth defects by *nusA11.* 10-fold serial dilutions of cultures with and without the *holC* complementing plasmid (pAM34-*holC*) were plated on minimal glucose casamino acids (min CAA) or LB media and incubated at the indicated temperature. These strains are derived from AB1157 (Rac^-^) in which lethal effects of *nusA* mutations are reduced.

To further explore the role of transcriptional termination in the phenotypes of *holC* mutants, we examined a mutation in the β subunit of RNAP *rpoB8* (Q513P) that increases transcriptional pausing, has a slower elongation speed and is more prone to Rho-dependent termination (Fisher and Yanofsky 1983; Landick et al. 1990; Jin and Gross 1991). The *rpoB8* allele significantly suppressed *holC* inviability, indicating that Rho-dependent termination aids the viability of *holC* mutants (Fig. 8). Interestingly, suppression was mutual. The *holC* mutation also suppressed the poor growth of *rpoB8* strains on either min CAA or LB; the plasmid-less *holC rpoB8* double mutant grew more robustly than either *holC* or *rpoB8* single mutants. We also examined effects of *rpoB3770* (T563P) that, like *rpoB8,* confers resistance to rifampicin. This is a “stringent” allele of *rpoB*, that suppresses phenotypes of mutants defective in mounting the stringent response to starvation via accumulation of the signaling molecule (p)ppGpp (Zhou and Jin 1998). In contrast to *rpoB8*, *rpoB3370* did not ameliorate *holC* growth phenotypes and further reduced colony size upon loss of the *holC*^+^ complementing plasmid. This finding is consistent with a *holC*-suppressive effect of Rho-dependent termination, since strains carrying *rpoB3370* exhibit decreased termination at three different Rho-dependent terminators from bacteriophage lambda (Jin et al. 1988).

**Figure 8.**
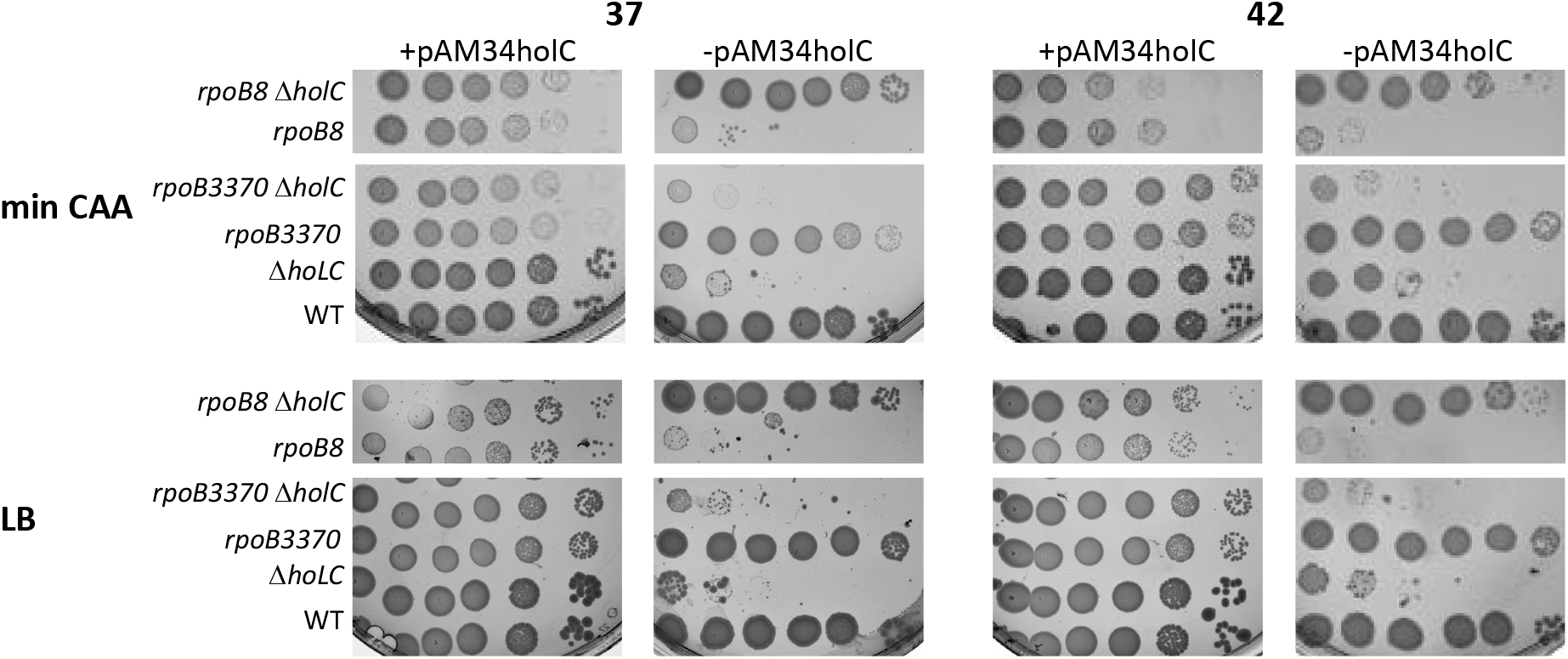
Suppression *holC* growth defects by *rpoB8,* enhancement by *rpoB3370.* 10-fold serial dilutions of cultures with and without the *holC* complementing plasmid were plated on minimal glucose casamino acids (min CAA) or LB media and incubated at the indicated temperature.

The antibiotic bicyclomycin is a specific inhibitor of Rho-dependent termination (Zwiefka et al. 1993). Treatment of *E. coli* cells with biocyclomycin induces replication-dependent double-strand breaks into DNA, indicative of the collapse of replication forks (Washburn and Gottesman 2011). Moreover, mutations that weaken transcription elongation complexes partially suppress this effect, supporting the hypothesis that Rho displaces RNAP before or after its collision with the replisome (Washburn and Gottesman 2011). We found that *holC* mutants were abnormally sensitive to the killing effects of bicyclomycin, consistent with the notion that transcription/ replication conflicts are more prevalent or more deleterious in the absence of HolC (Fig. 9AB). The effect is seen at both 30° and 37°, where plating efficiency of *holC* is reduced about 10-fold by 25 μg/ml BCM, whereas that of wild-type is unchanged. Loss of *sspA* completely suppresses *holC* at 30° on Min CAA medium; suppression is reduced 2 orders of magnitude by BCM (Figure 9A). Likewise, the *rpoAdup[aa179-186]* completely suppresses *holC* and suppression is abolished by BCM (Figure 9A). Neither *sspA* or *rpoAdup[aa179-186*) by themselves promote sensitivity to BCM (Figure 9A). Suppression of *holC* by *rpoC* is also lost in the presence of bicyclomycin (Figure 9B). This supports the notion that Rho-dependent termination, specifically inhibited by BCM, is required to sustain viability in the absence of *holC.* In addition, the ability of *rpoA, rpoC* and *sspA* mutations to suppress *holC* is dependent on Rho-dependent termination, indicating that these suppressor alleles may act through effects on Rho-dependent transcriptional termination.

**Figure 9.**
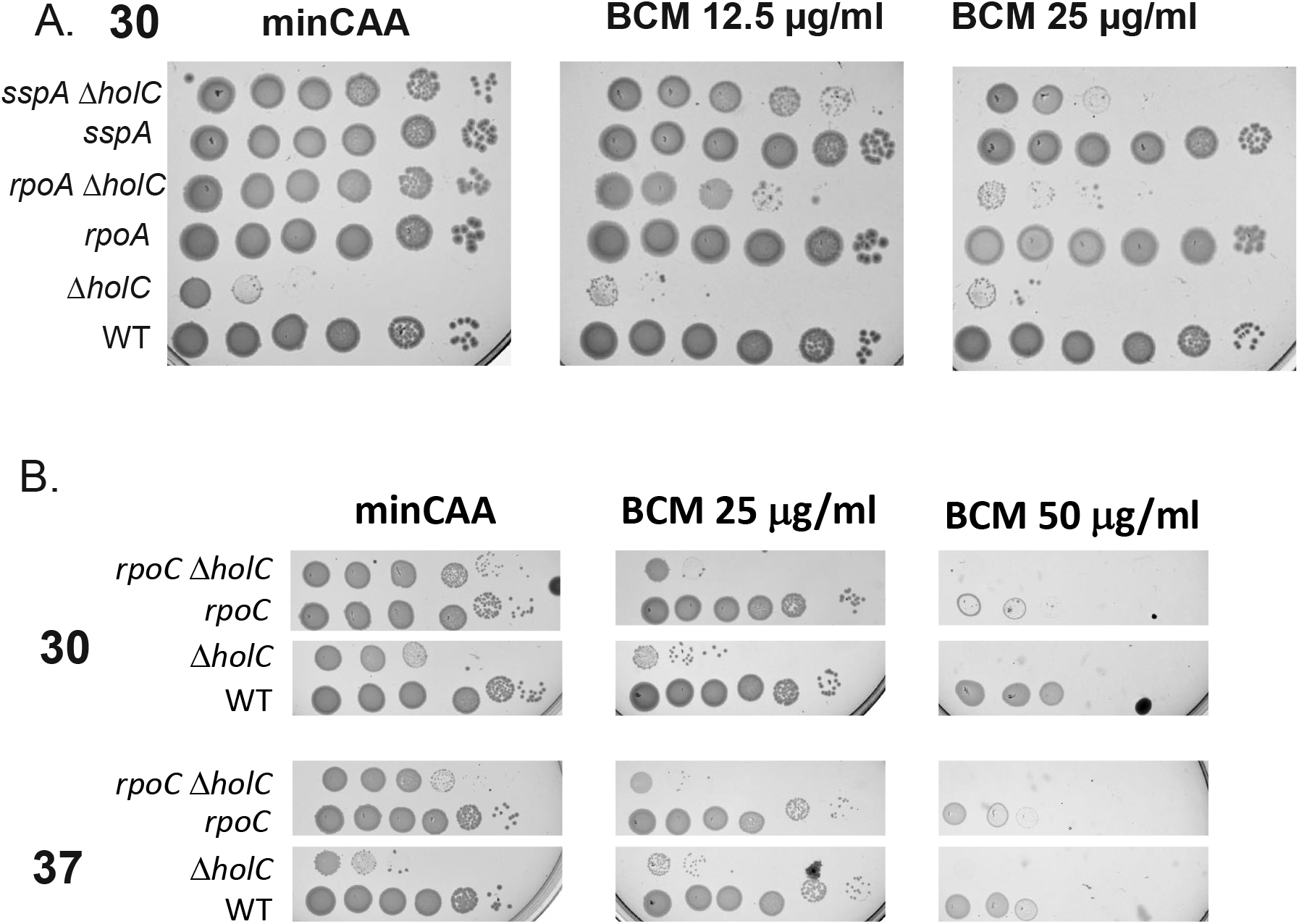
Bicyclomycin (BCM) sensitivity of *holC* mutants and *holC* suppression. 10-fold serial dilutions of cultures cured of the *holC* complementing plasmid were plated on minimal glucose casamino acids (min CAA) without and with two doses of bicyclomyin. A. The *rpoA* allele is *rpoAdup[aa 179-186]* and the *sspA* is a deletion. Suppression of *holC* by these alleles is reduced or abolished at 30° on min CAA medium. B. The *rpoC* allele is the *rpoC*::GFP allele, which partially suppresses *holC* on min CAA at 30° and 37° but not in the presence of bicyclomycin.

## DISCUSSION

Transcription elongation complexes are known to be impediments to the replication fork (reviewed in (Mirkin and Mirkin 2007; Merrikh et al. 2012; Garcia-Muse and Aguilera 2016)) and cells have evolved mechanisms to deal with these inevitable conflicts. Collisions between the replisome and transcription elongation complexes can occur in two orientations, head-on or codirectional (Fig. 11): of the two, head-on collisions are more deleterious, both in vivo (French 1992; Mirkin and Mirkin 2005; Mirkin et al. 2006) and in vitro (Liu and Alberts 1995; Pomerantz and O’Donnell 2008; Pomerantz and O’Donnell 2010). In bacterial genomes, gene orientation, especially for essential genes, is skewed so that most conflicts would be codirectional (Brewer 1988; Rocha and Danchin 2003). For example, in *E. coli* all 7 rRNA operons are arranged codirectionally with the fork. Reversing this orientation leads to transcription stalling, increased prevalence of RNA/DNA hybrids, and requirement for helicase proteins Rep, UvrD and DinG (Boubakri et al. 2010)). In *Bacillus,* inversion of a a locus is even more deleterious, leading to growth impairment even in the presence of analogous helicases (Wang et al. 2007; Srivatsan et al. 2010).

The absence of HolC perturbs DNA replication in several ways. The suppression of *holC* by duplication of the *ssb* gene (Duigou et al. 2014) suggests that replication is incomplete and the chromosome accumulates ssDNA gaps. In vitro, DNA replication with HolC mutants defective in SSB binding leads to uncoupling of leading and lagging strand synthesis with poor leading strand synthesis (Marceau et al. 2011). The in vivo results presented here suggest that another function of HolC protein may be to overcome or avoid replication conflicts with transcription elongation complexes. A mutation known to reduce the stability of RNAP, RpoA R191C (Kawakami and Ishihama 1980; Igarashi et al. 1990), was isolated as a suppressor of the poor growth phenotype exhibited by *holC* mutants. Additional mutations in RpoA (α), RpoB (β) and RpoC (β’) subunits of RNA polymerase also acted as suppressors of *holC;* although it is unclear what biochemical defects are caused by these alleles, we think it is likely that they represent some loss of function in RNAP, Conversely, transcription factor DksA and Rho-termination factor NusA sustain viability in the absence of HolC and loss of their functions leads to synthetic growth defects with *holC.* DksA has been best studied for its role in regulation of transcriptional initiation, where it potentiates the effects of the stringent response signaling molecule, ppGpp, on RNAP; in vivo it is required to downregulate rRNA synthesis during amino acid starvation (Gourse et al. 2018). *E. coli dksA* mutants are more prone to replication stalling and induction of the SOS response after amino acid starvation, in a manner that is reversed by inhibition of transcription with rifampicin (Tehranchi et al. 2010). DksA is also required for replication initiated by RNA/DNA hybrids (“R-loops) and may also assist in the removal of RNAP (Myka et al. 2019). NusA mutants are hyper-sensitive to bicyclomycin, an inhibitor of Rho-dependent termination, and exhibit more chromosomal fragmentation during replication (Washburn and Gottesman 2011), implicating a role for Rho-dependent termination in sustaining chromosome integrity. This work extends this finding and shows that Rho-dependent termination must be particularly critical in the absence of HolC, potentially to clear transcription elongation complexes to avoid collision with the replication fork. How HolC’s two binding partners, HolD (the clamp loader protein) or YoaA (putative helicase), participate in this role remains to be determined.

The factors required to mitigate transcription/replication collisions are complex and potentially situation-specific. In *E. coli,*YoaA has a paralog, DinG, which is one of the DNA helicases required to survive head-on replication/transcription collisions in highly expressed genes (Boubakri et al. 2010). It is tempting to speculate that HolC/YoaA may aid tolerance of co-directional replication/transcription collisions, as would occur at *rrn.* Although paralogous, DinG and YoaA appear to have distinct functions: *yoaA* but not *dinG* confers sensitivity to AZT when deleted and resistance when overexpressed (Brown et al. 2015), indicating they are not merely redundant and must have specialized roles.

Because DnaB, the replication fork helicase, translocates on the lagging-strand template, a codirectional collision of the replisome with RNAP elongation complexes leads to different outcomes than a head-on collision (Fig. 10). In the codirectional orientation, DnaB can proceed unimpeded, uncoupling leading and lagging strand synthesis. The codirectional orientation can potentially lead to the use of RNA component of a RNA/DNA hybrid (or “R-loop”) to reprime DNA synthesis. It has been documented that DksA aids in the use of R-loops to initiate DNA synthesis and may assist in removal of transcription elongation complexes to facilitate repair, which may explain how DksA sustains growth in *holC* mutants.

**Fig. 10.**
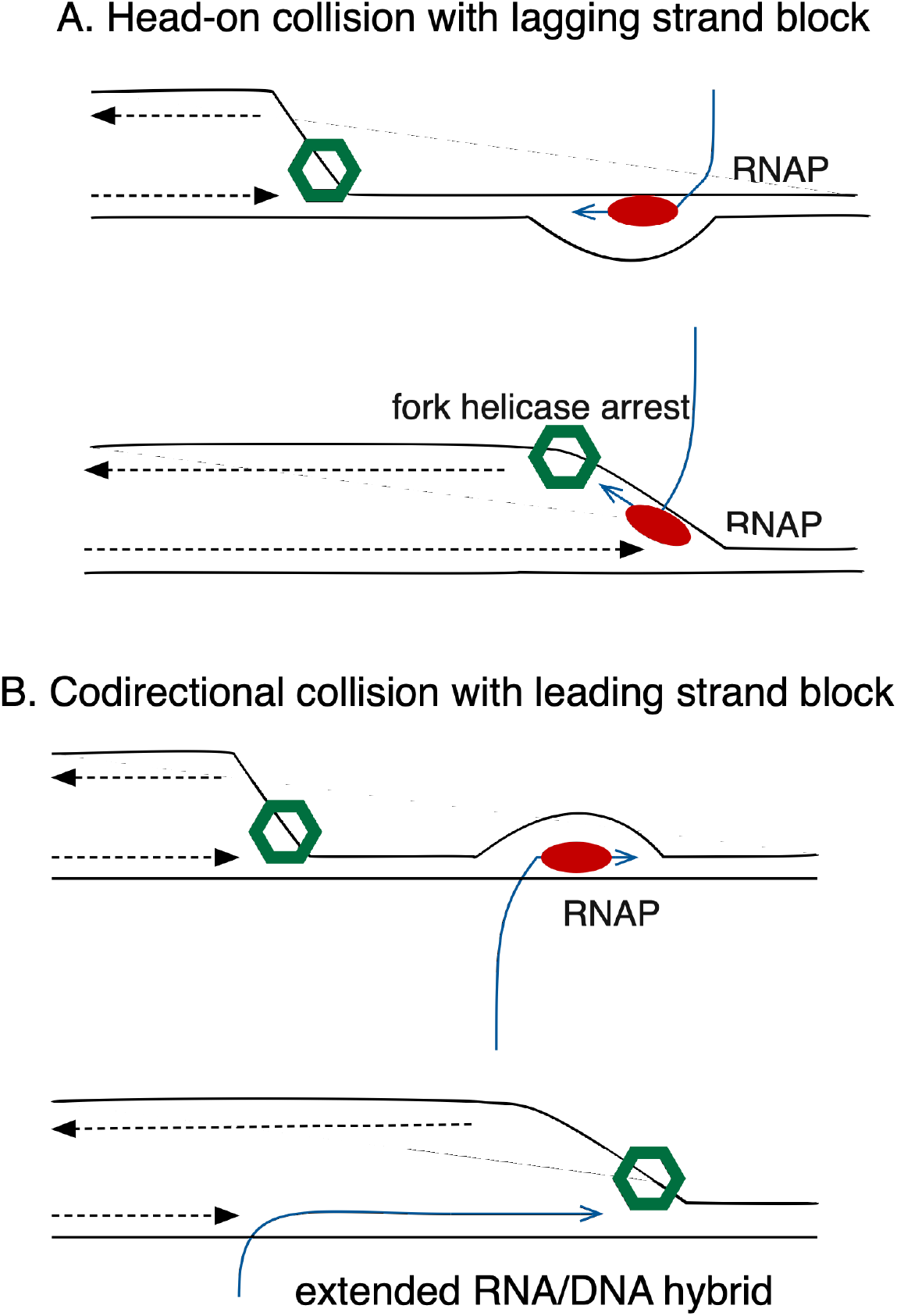
Transcription/replication conflicts. In green is illustrated the DnaB fork helicase; in red RNA polymerase, DNA in black; RNA in blue. A. Head-on collisions lead to fork arrest B. Codirectional collisions cause uncoupling of leading and lagging strand synthesis and possible stabilization of R-loops.

In addition to alleles of RNAP, we also isolated alleles of *sspA* as growth suppressors of *holC.* Mutations is *sspA* have been shown previously to suppress loss of *holD* (Michel and Sinha 2017). SspA, “stringent starvation protein”, is a growth-regulated RNA polymerase-associated protein (Ishihama and Saitoh 1979; Williams et al. 1994a), that can act as an activator of gene expression (Hansen et al. 2003). Although it is primarily expressed during stationary phase of growth, it also regulates, either directly or indirectly, a number of genes during exponential growth (Williams et al. 1994b). SspA promotes replication of bacteriophage P1(Hansen et al. 2003) as well as resistance to acid stress (Hansen et al. 2005), long-term starvation (Williams et al. 1994b) and is required for virulence for many bacterial pathogens (De Reuse and Taha 1997;

Badger and Miller 1998; Baron and Nano 1998; Tsolis et al. 2000; Merrell et al. 2002; Hansen and Jin 2012; Honn et al. 2012). In *E. coli,* it down-regulates nucleoid-associated protein H-NS (Hansen et al. 2005; Hansen and Jin 2012). However, the suppressive effect of *sspA* on *holD* does not appear to be due to increased H-NS, since H-NS overexpression by itself does not improve the viability of HolD (Michel and Sinha 2017).

Whatever its mechanism, suppression of *holC* is likely to be similar to *holD* but the downstream effector(s) or mechanism of SspA responsible for this suppression is currently unknown. Our observation that suppression of *holC* by *sspA* is negated in the presence of bicyclomycin suggests that it may act by affecting termination or RNAP properties, either directly or indirectly. Given that SspA is a transcriptional activator, SspA may induce something deleterious to *holC* mutant strains, although it is possible that something advantageous to *holC* mutants is repressed or RNAP properties are altered in a more general way. The potential links between SspA and DNA metabolism will require further study.

## MATERIALS AND METHODS

**Table 1**. All alleles for mutations are derived from the Keio collection (Baba et al. 2006) except as noted. All strains listed except the wild-type strains AB1157 (STL140), MG1655 (Bachmann 1996) and PFM2 (MG1655 rph+)(Lee et al. 2012) have been transformed with the pAM34-holC plasmid. The construction of the pAM34-holC plasmid is described in the materials and methods.

**Table 1.**
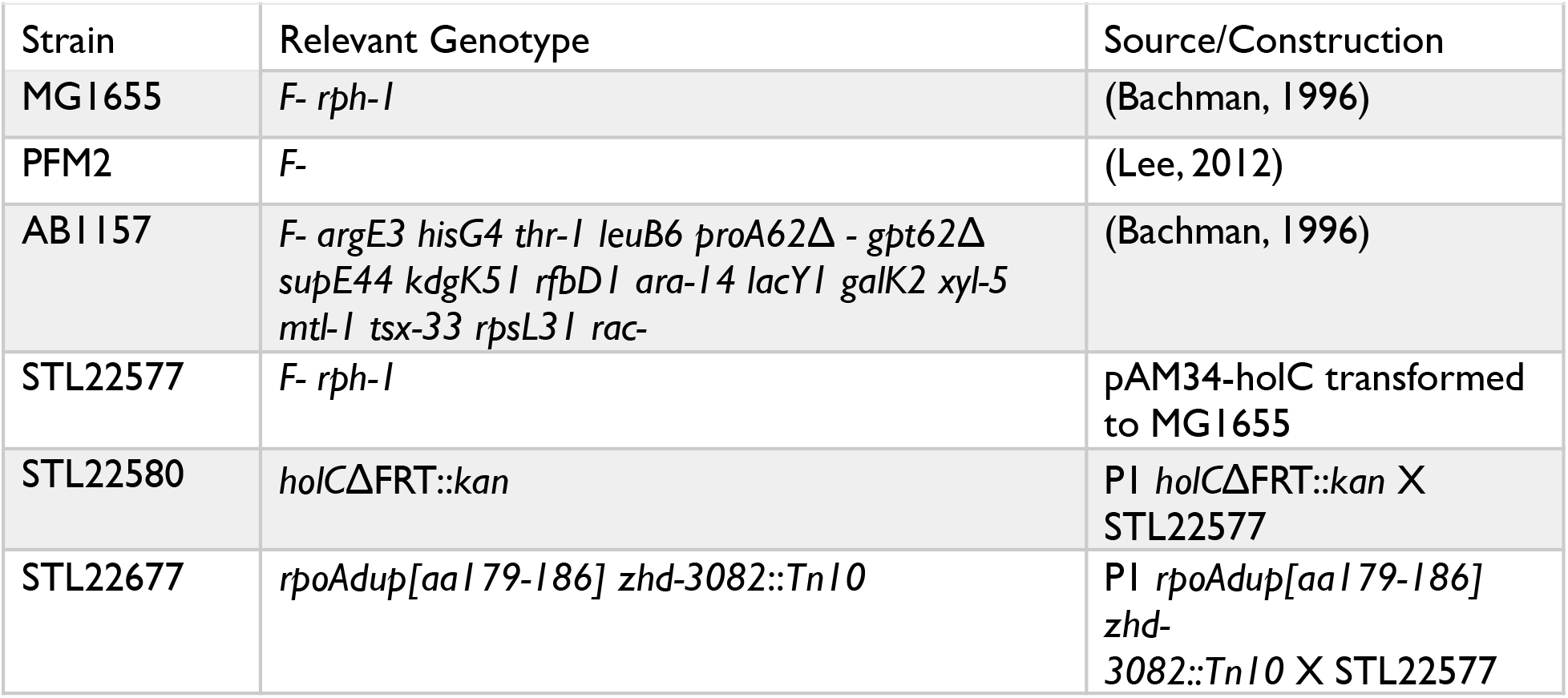
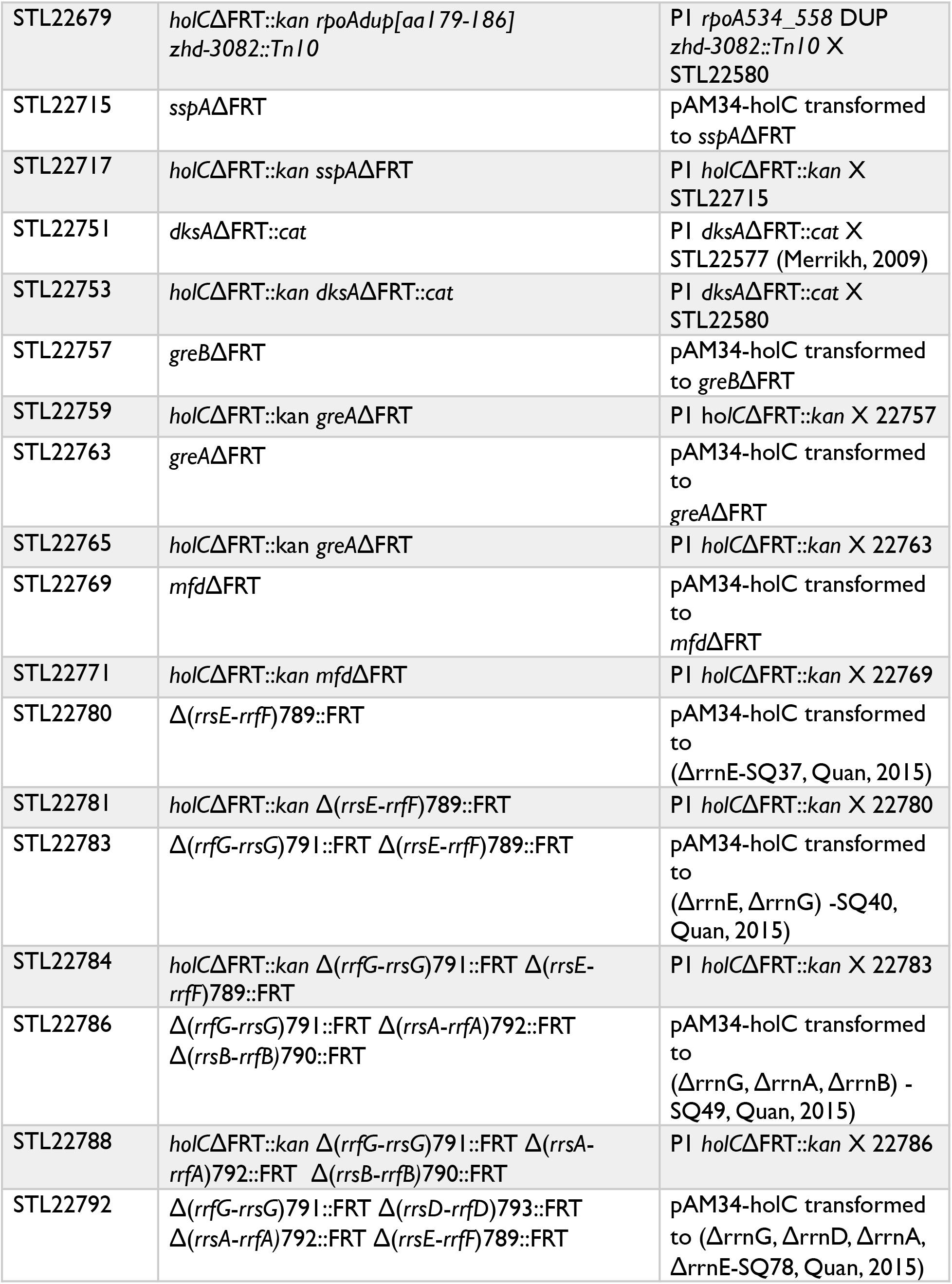
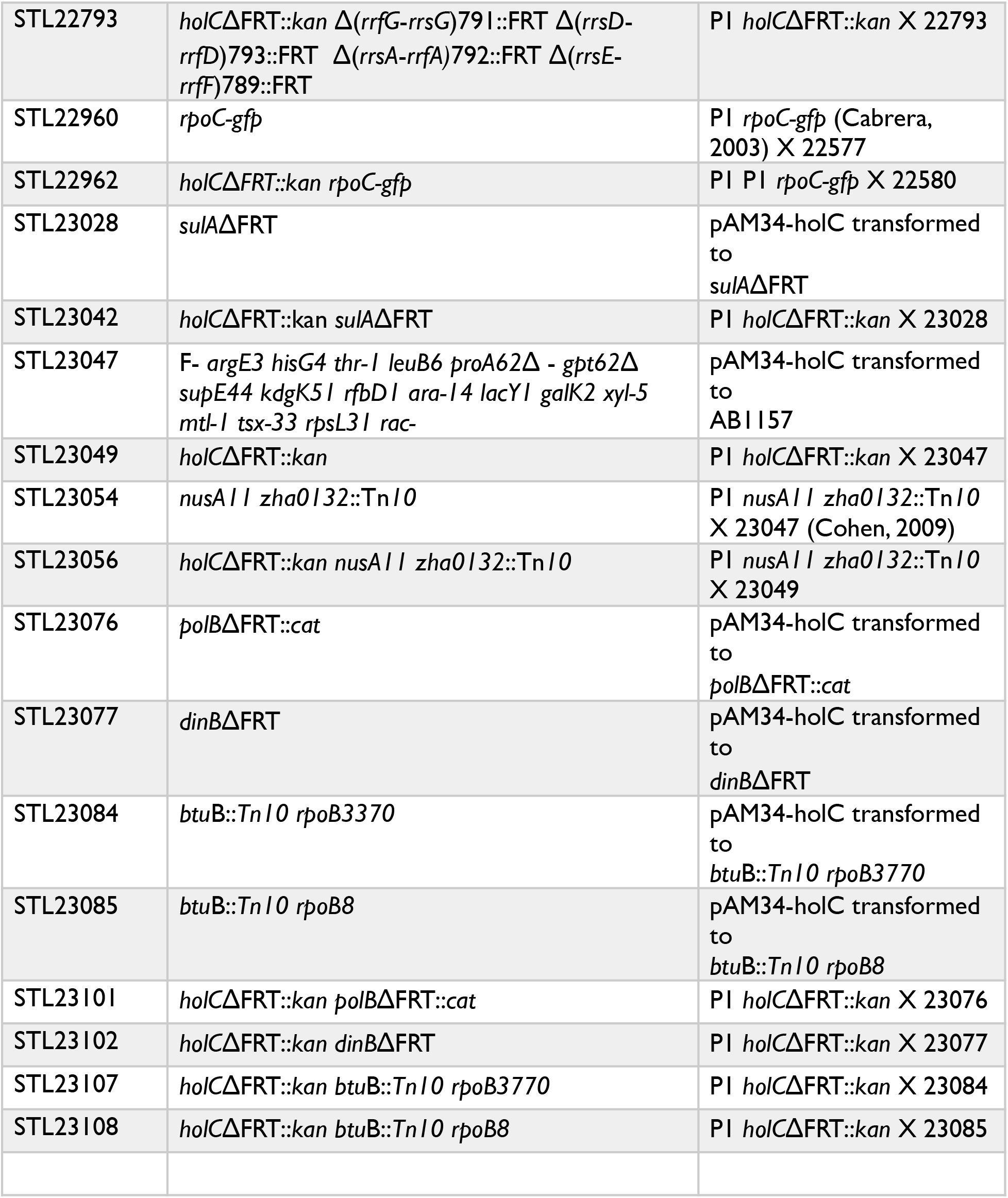
*E.coli* K-I2 strains and plasmids

### Routine Growth

Bacterial cultures were routinely grown at 30° in minimal media + casamino acids (Min-CAA) containing 56/2 salts, 0.2% (w/v) glucose, (0.001% (w/v) thiamine, and 0.2% (w/ v) CAA. Plate media contained 2.0% (w/v) agar. For experiments testing the effects of media on growth, LB media or minimal media were also used. LB media contained 1% (w/v) tryptone, 0.5% (w/v) yeast extract, 0.5% (w/v) NaCl with 1.5% (w/v) agar for plates. Minimal media (Min) was identical to Min-CAA except the CAA were omitted.

### Strain Construction

The Escherichia coli strains used here were all MG1655 derivatives, except for the strains containing *nusA11* which were AB1157 derivatives. All alleles for mutations were derived from the Keio collection (Baba et al. 2006) except *dksA* (Merrikh et al. 2009), *rrn* (Quan et al. 2005), *nusA11* (Cohen et al. 2009), *rpoB8* (Washburn and Gottesman 2011), *rpoC*::GFP (Cabrera et al. 2009), *rpoB3370* (Jin and Gross 1989). To decrease the possibility of unintended suppressors arising, strains also contained a pAM34-*holC* plasmid which allows conditional expression of *holC*.

Strains were constructed using P1 transduction as described (Miller 1992). HolC was expressed from pAM34-*holC* during all strain preparation involving a *holC* deletion allele. For preparation of P1 lysates on *holC* deletion strains or the *rpoAdup*[aa179-186] strain, minimal media with 2mM CaCl2 was used. Plates contained 1.5% agar and top agar contained 0.7% agar. This same media was also used for transduction into these strains. For all other strains, LCG media was used for preparation of P1 lysates and transductions. LCG consisted of LB media supplemented with 0.1% (w/v) glucose and 2 mM calcium chloride with plate medium containing 1% (w/v) agar and LCG top agar containing 0.7% (w/v) agar. Transductants were selected at 30° on Min-CAA plates containing the appropriate antibiotic and 0.15-0.2 mM IPTG and ampicillin when the pAM34-*holC* plasmid was present. The following antibiotic concentrations were used: 100 μg/ml of ampicillin (Ap), 30 μg/ml of chloramphenicol (Cm), 60 μg/ml of kanamycin (Km) and 15 μg/ml of tetracycline (Tc). Bicyclomycin (BCM) was kindly provided by Robert Washburn. Strain constructions were verified by phenotype, PCR and/or sequencing.

### Plasmid Construction

The pAM-34-*holC* plasmid was constructed from pAM34 which was kindly provided by Bénédicte Michel. The *Xöa*I-*Sac*I fragment of pAM34 which contains the spectinomycin gene was replaced with a DNA fragment containing *holC* and its 100 bp upstream region to allow expression of *holC* from its natural promoter. Replication of this plasmid depends on IPTG. In the experiments described here, IPTG was added to between 0.15 and 0.2 mM to help maintain a low plasmid copy number and minimize deleterious effects of *holC* overexpression on cell growth.

### Growth Experiments

To test the growth of *holC* mutants, strains were grown from single colonies in the presence of 0.15-0.2 mM IPTG and ampicillin for 10-12 hours in min-CAA media at 30°. Cultures were then split and diluted to an A_590_ of approximately 0.005 in either min-CAA media containing ampicillin and IPTG (pAM34-holC maintained) or in min-CAA media alone (pAM34-holC lost). Growth was continued for 14-16 hours. Next, cultures were diluted into the same media and allowed to grow to mid-late log phase (6-8 hours) at which time they were serially diluted and plated on LB, min-CAA, and min plates at 30, 37, and 42 as indicated in the figure legends. All experiments were done with multiple biological isolates and repeated on at least two days except as noted in the figure legends.

### Microscopy

Cells depleted of pAM34-holC were fixed by adding an equal volume of methanol:acetic acid (3:1) to the liquid cultures. Fixed cells were then spotted onto poly-L-Lysine treated slides, washed extensively with PBS, and overlaid with Vectashield Mounting Media. Slides were then imaged using phase contrast with an Olympus BX51 microscope and a Qimaging Retiga 559Exi camera. The cell lengths of all of the cells in any given field of view was determined using ImageJ (Abramoff 2004).

## Acknowledgments

We thank Bénédicte Michel for providing the pAM plasmid vector, Graham Walker for the *nusA11* strain, Robert Washburn for the *rpoB8* strain and for bicyclomycin, Cathy Squires for the *rrn* deletion strains, Ding Jin for *rpoC*::GFP and *rpoB3370* strains and the National Genetic Institute of Japan for the Mori knockout collection. This work was supported by NIH grant R01 GM51753. We thank Rick Gourse and Mike Marr for discussions regarding transcription and its termination.

---

Supplemental Figure 1. Lack of effect on *holC* growth defects by *greA and mfd.* 10-fold serial dilutions of cultures with and without the *holC* complementing plasmid were plated on minimal glucose media and incubated at the indicated temperature.

## Footnotes

1 Because the same Greek letters are used for subunits of DNA polymerase III and RNA polymerase, for simplicity we use gene names here to designate the DNA pol III proteins and Greek letters for RNAP.

