## Supplemental Figure 1 for "The role of replication clamp-loader protein HolC of *Escherichia coli i*n overcoming replication / transcription conflicts"

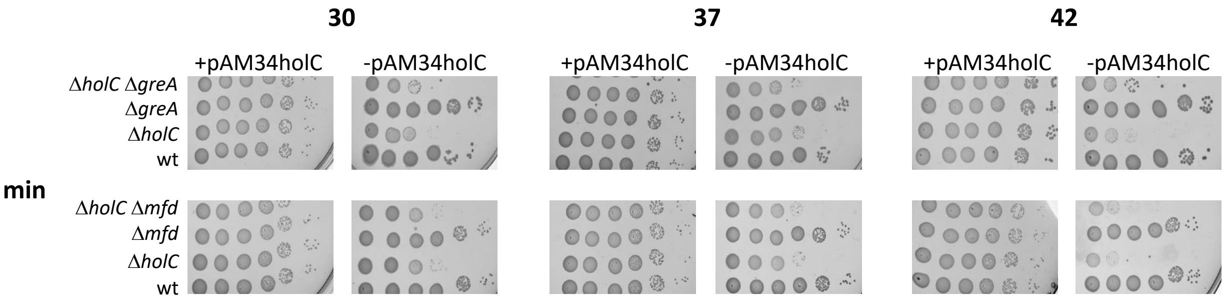


Supplemental Figure 1. Lack of effect on *holC* growth defects by *greA and mfd.* 10-fold serial dilutions of cultures with and without the *holC* complementing plasmid were plated on minimal glucose media and incubated at the indicated temperature.
